# Orchestration of Proteins in cyanobacterial Circadian Clock System 1

**DOI:** 10.1101/2022.08.26.505376

**Authors:** Masaaki Sugiyama, Ken Morishima, Yasuhiro Yunoki, Rintaro Inoue, Nobuhiro Sato, Hirokazu Yagi, Koichi Kato

## Abstract

Circadian rhythm by *Cyanobacteria* is one of the simplest biological clocks: the clock consists of only three proteins, KaiA, KaiB, and KaiC. Their oligomers, KaiA dimer (A_2_), KaiB tetramer (B_4_), and KaiC hexamer (C_6_) oscillate an association–disassociation cycle with 24-hr period. In a widely accepted model, the oscillation process is as follows. From the viewpoint of a base unit (C_6_), C_6_ homo-oligomer → A_2_C_6_ complex → B_6_C_6_ complex → A*_n_*B_6_C_6_ complex (*n*≤12) →C_6_ homo-oligomer. In this study, Small-Angle X-ray Scattering, Contrast Matching-Small-Angle Neutron Scattering, Analytical Ultracentrifuge, and phosphorylation-analysis PAGE measurements were performed to reveal the kinetics not only of KaiC hexamer but also of all components in a working Kai clock. The complementary analysis disclosed that the oscillation is not the single process as the widely accepted model but composed with synchronized multiple association-dissociation reactions between components. Namely, there are various reactions between components, which proceed simultaneously, in a working Kai-clock.

## INTRODUCTION

Every life on this planet has a circadian clock to regulate the metabolic rhythm along with a daily circle. Cyanobacteria has one of the simplest circadian clocks which consists of only three proteins, KaiA, KaiB and KaiC, named as Kai clock [1]. Kai clock works even *in vitro* solution with adenosine tri-phosphate (ATP) but without any daylight oscillation [2, 3]. Therefore, Kai clock has been intensively investigated as one of model systems in the last decades [4, 5, 6].

A Kai clock system includes two rhythms in the different scales, intra-molecular and inter-molecular ones. The intra-molecular rhythm is an oscillation of a phosphorylation degree of a KaiC molecule (PhosD) [7]. A KaiC molecule has two phosphorylation sites, Ser431 and Thr432, of which the phosphorylation oscillates with 24-hr period under a regularly-working clock as follows, ST→S*p*T→*p*S*p*T→*p*ST→ST (*p* indicates a phosphorylated residue.) [8]. Considering that a KaiC molecule has dephosphorylation nature, the Kai clock system should have the KaiC-phosphorylation mechanism which is activated in the first two processes (ST→S*p*T→*p*S*p*T) and is inactivated in the last two ones (*p*S*p*T→*p*ST→ST). Furthermore, the phosphorylation mechanism simultaneously works for a majority of KaiC molecules in the clock system because the averaged PhosD over the KaiC molecules in the system also oscillates with 24-hr period [9]. Therefore, it is rational that there should be an inter-molecular rhythm which regulates the KaiC-phosphorylation mechanism.

The inter-molecular rhythm is an oscillation of complex formation and the subsequent deformation between three homo-oligomers, Kai A dimer (A_2_) [10], KaiB tetramer (B_4_) [11] and KaiC hexamer (C_6_) in a solution with ATP [12]. They and their complexes have particular features correlating with the phosphorylation ability for a KaiC molecule as follows. A free A_2_ associates with a dephosphorylated C_6_, generating A_2_C_6_ complex. The A_2_C_6_ phosphorylates its KaiC protomers and then the A_2_C_6_ with the phosphorylated C_6_ protomer is likely decomposed into A_2_ and C_6_ oligomers, again [13]. Then, a free B_4_ associates with the phosphorylated C_6_, generating B_6_C_6_ complex, in the phosphorylation phase around *p*S*p*T→*p*ST [14, 15]. Subsequently, the B_6_C_6_ associates with a free A_2_ to produce A*x*B_6_C_6_ (*x* is supposed to reach 12) [16, 17]. A*x*B_6_C_6_ has no phosphorylation ability and then the A*x*B_6_C_6_ with the dephosphorylated C_6_ protomer is likely decomposed into A_2_, B_4_ and C_6_ oligomers, again.

The widely accepted picture of the oscillation process has been presented by coupling with the intra- and inter-molecules rhythms [16]. As shown in Fig.S1, from viewpoint of C_6_ hexamer, in the starting point (Point a), C_6_ is homo-oligomer, meaning that this is a fully decomposed state, and then, in Point b, C_6_ forms A_2_C_6_ complex by Process i and is phosphorylation-activated, and then, in Point c, C_6_ forms B_6_C_6_ complex, by Process ii and is phosphorylation-inactivation [18]. Subsequently, in Point d, the A*x*B_6_C_6_ (*x* → 12) is formed by Process iii, which induces the shortage of free A_2_ oligomers. Therefore, the formation of ABC complex suppresses the formation of A_2_C_6_ (Process i) and then the formation of B_6_C_6_ (Process ii) is also suppressed. As a result, the components in the system are concentrated to be A*x*B_6_C_6_ complexes (*x* → 12). This is supposed to be a phase-synchronized mechanism. Then, KaiC in A*x*B_6_C_6_ (*x* → 12) is dephosphorylated, subsequently A*x*B_6_C_6_ (*x* → 12) is decomposed to the homo-oligomers, A_2_ B_6_ and the dephosphorylated C_6_ (Process iv). Then, the system returns the starting point (Point a).

To examine the above-mentioned picture, we measured a working Kai clock with time-resolved Small-Angle X-ray Scattering (SAXS) and phosphorylation-analysis-PAGE (see Materials and methods). Figure S2 shows the time evolutions of the forward SAXS intensity *I*(0) and the averaged PhosD. The observed *I*(0) and the averaged PhosD oscillated with a 24 hr-period, which well-reproduced the previous report [18]. To consider the relation between these oscillations and the widely-accepted picture, based on the time evolutions of PhosD, we divided the measured 24 hrs into four process zones as shown with horizontal black arrows marked by numbers as P I, P II, P III and P IV: Process I (P I) is a phosphorylation-acceleration zone, and Process II (P II) is a phosphorylation-deceleration zone, and Process III (P III) is a dephosphorylation-acceleration zone, and Process III (P III) is a dephosphorylation-deceleration zone. Here, we noticed two boundary time points, t_min_ (24 hr) and t_max_ (36 hr), where the *I*(0) values reached the minimum and maximum values, *I*_min_(0) ≅ 0.65 cm^-1^ and *I*_max_(0) ≅ 0.82 cm^-1^, respectively pointed by arrows in Fig.S2. Following the widely accepted picture, we considered the molecular compositions at these points. At t_min_, the first candidate of molecular composition could be a state fully-decomposed to homooligomers (comp 1: A_2_, B_4_ and C_6_ in the system, namely Point a) because the *I*(0) value was minimum. The *I*(0) value of comp 1, *I*_1_(0), is calculated to be 0.458 cm^-1^ indicated by black broken line in Fig.S2: *I*_1_(0) is reasonable coincident with the *I*(0) value at 0 hr but *I*_1_(0) > *I*_min_(0). This means that the larger oligomers, such as A_2_C_6_ and/or B_6_C_6_, could be coexisted at t_min_. Following this idea, another two compositions were simulated as the upper limit *I*(0) values at t_min_: one (comp 2) is a combination of A_2_C_6_, B_4_ and remaining C_6_, where KaiA molecules are fully associated to C_6_ and the other (comp 3) is a combination of B_6_C_6_, A_2_ and remaining B_4_, where KaiB molecules are fully associated to C_6_: The comp 2 and comp3 could correspond to Points b and c, respectively. As shown in Fig.S2, the calculated *I*(0) values of comps 2 and 3, are *I*_2_(0) = 0.619 cm^-1^ (blue broken line) and *I*_3_(0) = 0.612 cm^-1^ (green broken line), respectively. It should be stressed that these values are still lower than *I*_min_(0). Accordingly, there could be not only A_2_C_6_ and /or B_6_C_6_ but the larger complexes such as ABC complex even at t_min_. Namely, in a working Kai clock, there is not simple molecular distribution such as that of Point b or c. Likewise, the time point at t_max_ was supposed there could be the fully ABC complex (Point d) because the *I*(0) value became maximum. Therefore, it was supposed to that the molecular composition could be A12B_6_C_6_, B_6_C_6_ and B_4_ (comp 4). The calculated *I*(0) for comp 4 was *I*_4_(0) = 0.959 cm^-1^ (a red broken line in Fig.S2), which was much larger than *I*_max_(0). This means that comp 4 (pure Point d) does not realize in the real Kai clock system. There is no directly-connected time point nor processes between the widely-accepted picture and the experimental results, which reflected the molecular distributions in a whole Kai clock system.

In consideration of the above, all C_6_ oligomers do not synchronously transit the same pathway as shown in Fig.S1. On the other hand, all C_6_ oligomer does not randomly progress the oscillation pathway because the clock system exhibits the oscillation of forward SAXS intensity which represents the distribution of components. Therefore, there should be overlooked synchronous association and dissociation processes between the components in the Kai clock system. To access these processes experimentally, it is inevitable to reveal the kinetics of all components over the oscillation period. For this purpose, we measured the distribution of components in the working Kai clock by complementary use of time-resolved measurements, Analytical Ultracentrifuge (AUC), SAXS, Contrast Matching Small-Angle Neutron Scattering (CM-SANS) [19], and phosphorylation-analysis PAGE. The results tell us that the widely accepted picture describes only a part of Kai clock system.

## RESULTS

### Overview of time evolution of oscillation system

KaiA, KaiB and KaiC solutions with [KaiA]:[KaiB]:[KaiC] = 2.8: 7.8: 6.0 were mixed. Then, the AUC measurement was performed every two hours from 24 hr at the first forward *I*_min_(0) time point to 48 hr at the next *I*_min_(0) time point coupled with the time resolved SAXS and the KaiC-phosphorylation degree (see details in Materials and methods). Figure1 (a)-(l) shows the time evolution of the AUC profile, which are the weight concentration distributions of particles as a function of svedberg unit *c*(*s*_20,*w*_). Considering that the peak position and area correspond to the particle molecular weight and its number, the result clearly indicates that there is the time-transitional association-dissociation between the components in a working Kai clock. In addition, the AUC profile at 48 hr (a red line in Fig.1(b)) returned to that at 24 hr (a black line in Fig.1(b)), indicating that the association-dissociation oscillates with 24hrs period. To overlook the kinetics of all components, Figure 2 displays the time evolution of AUC spectra with twodimensional heat map as *x, y*, and *z* axes are an elapsed time after the mixture with the arrows indicating Processes I-IV, *s*-value (*s*_20,*w*_) and the weight concentration (*c*(*s*_20,*w*_)), respectively: Above the heat map, the time resolved *I*(0) and the PhosD are displayed as references. Surprisingly, three distinct streams, Stream 1 (S1), Stream 2 (S2) and Stream 3 (S3) in the order of *s*-value, are clearly recognized along the time course in the figure: The time evolution of their peak positions and intensities, *s-* and *c*-values, are also shown in Fig.3 (a) and (b), respectively. The features of three streams are summarized as follows.

**Figure 1.**
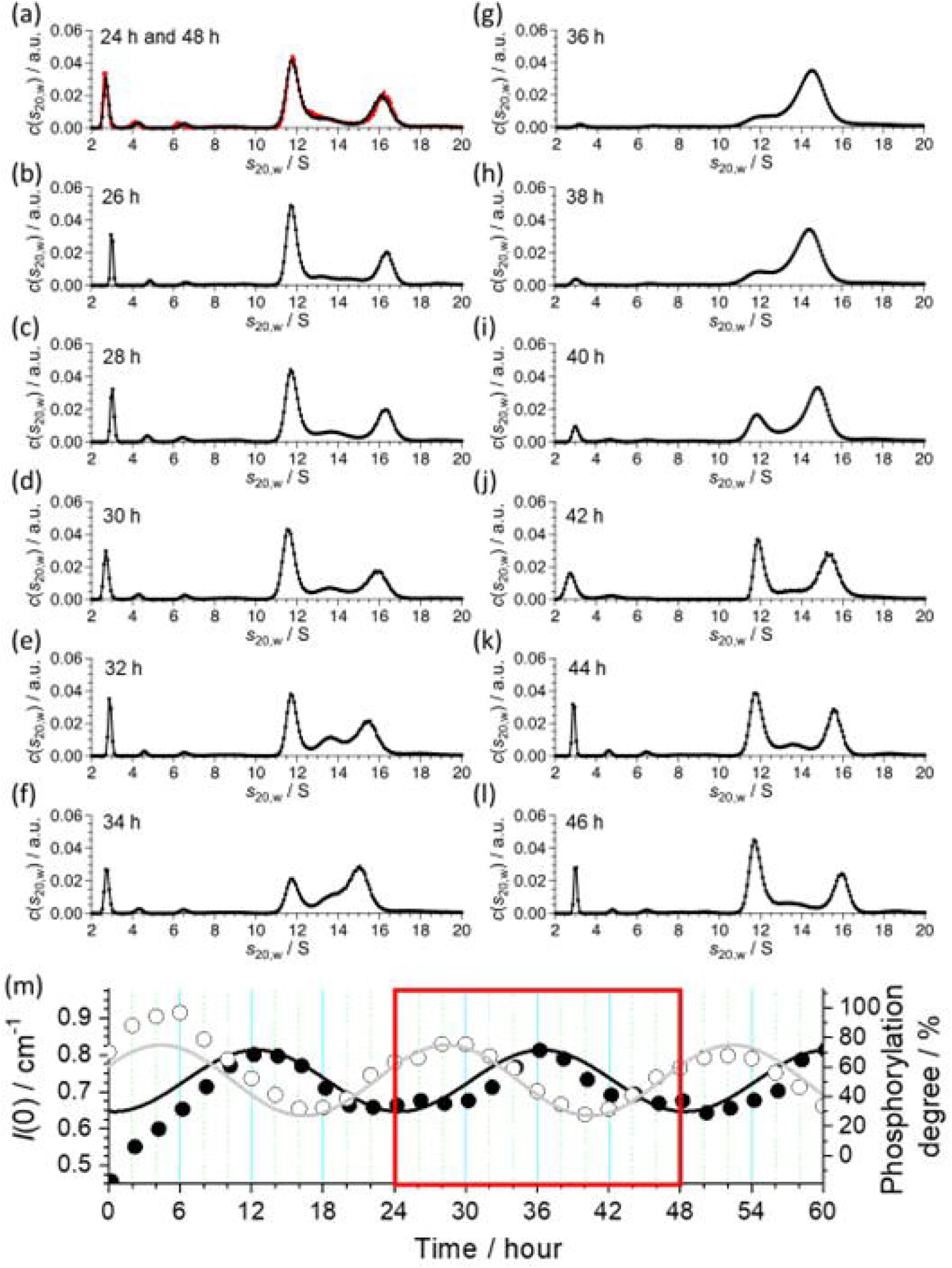
Time evolutions of AUC profiles and simultaneously-measured time evolutions of forward SAXS intensity and phosphorylation degree of KaiC. Time evolutions of AUC profiles at (a) 24hr (a black line) and 48hr (a red line), (b) 26hr, (c) 28hr, (d) 30hr, (e) 32hr, (f) 34hr, (g) 36hr, (h) 38hr, (i) 40hr, (j) 42hr, (k) 44hr, and (l) 46hr. (m) Simultaneously-measured time evolutions of forward SAXS intensity and phosphorylation degree of KaiC (PhosD). Close and open circles denote forward SAXS intensity and phosphorylation degree of KaiC, respectively. Black and grey lines are eye guide ones with a 24hrs-oscillation period. A red circle indicates the time zone where the AUC measurement was performed (See the detail in Materials and methods.).

**Figure 2.**
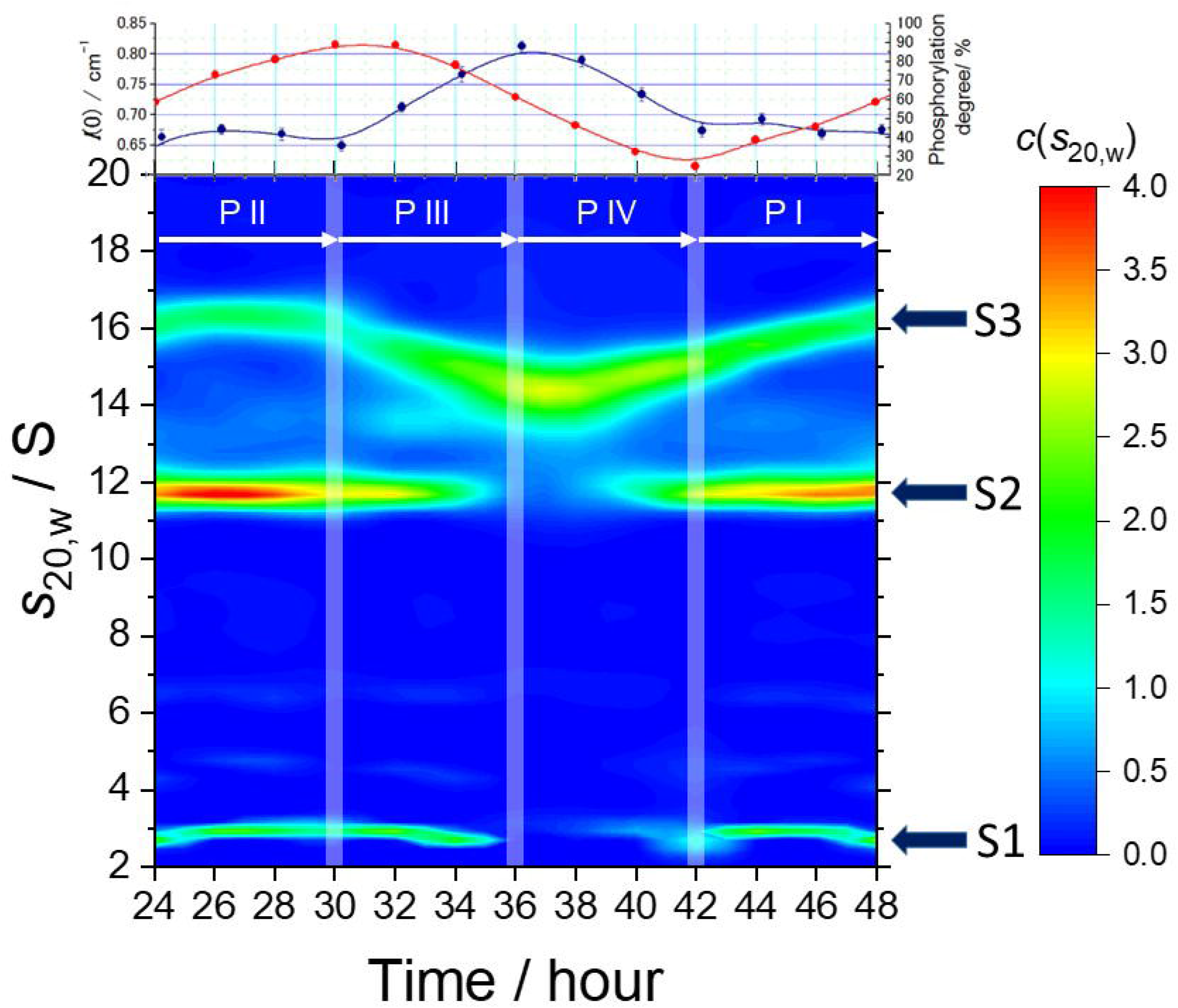
Two-dimensional map of time evolution of AUC-spectra: *x, y*, and *z* axes are an elapsed time after the mixture, *s*-value (*s*_20,*w*_), and the weight concentration (c(*s*_20,*w*_)), respectively. Thick dark blue arrows indicate three streams, Steam 1 (S1), Stream 2 (S2), and Stream 3 (S3) in the order of *s*-value. White arrows and thick half transparent white lines denote the time range of processes shown in Fig.S2 and their boundary, respectively. The upper panel shows the simultaneously measured forward SAXS intensity (blue) and phosphorylation degree of KaiC (red).

**Figure 3.**
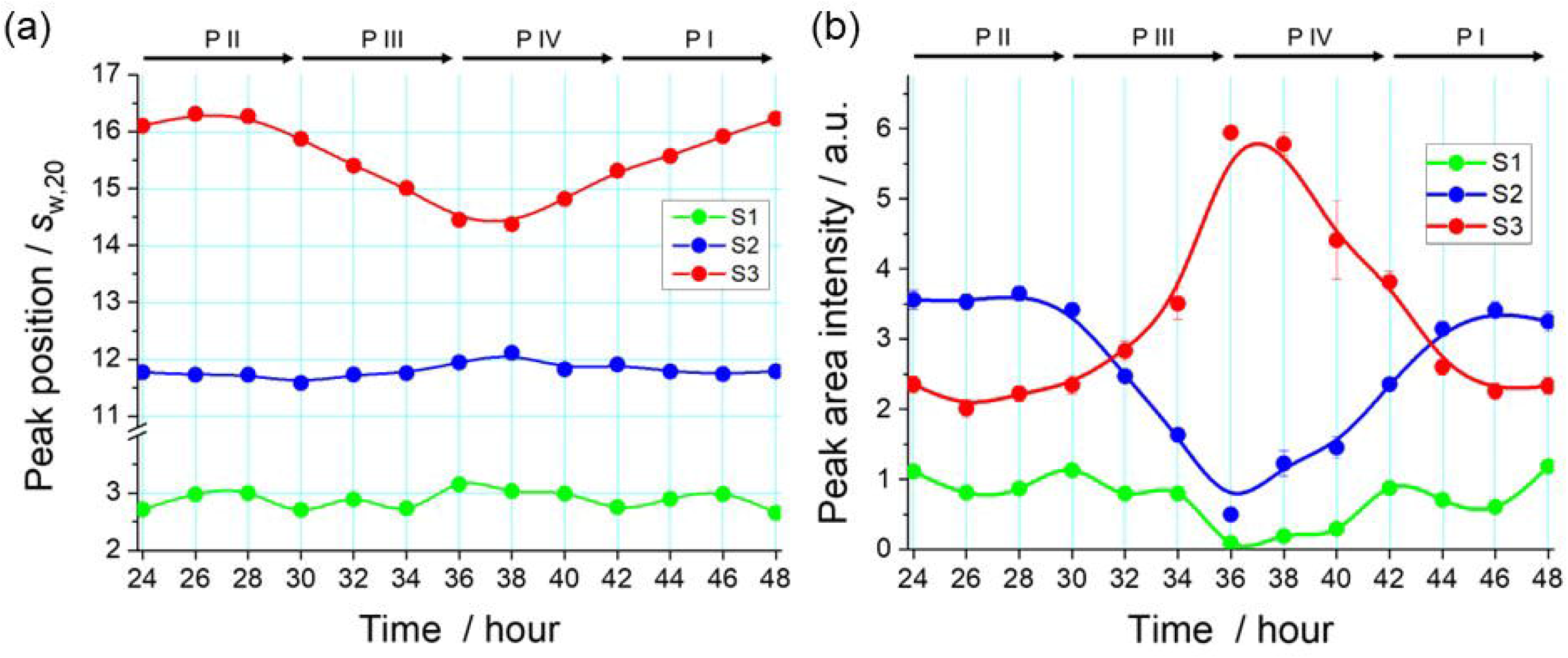
Time evolutions of peak positions and their areas of three streams in AUC profiles. (a) Green, blue and red circles indicate time evolutions of peak positions of Streams 1, 2, and 3, respectively. (b) Green, blue and red circles indicate time evolutions of peak areas of Streams 1, 2, and 3, respectively. In both panels, lines are eye guides.

#### Stream 1

The *s*-value was almost constant (2.83 ± 0.13S) whereas the *c*-values became close to zero between 36 hr and 40 hr (around a boundary between Processes III and IV). The later means that the all components corresponding to Stream 1 connected with the other component(s).

#### Stream 2

The *s*-value was almost constant (11.75 ± 0.08S) whereas the *c*-values oscillated with 24 hrs period. The *c*-values decreased from 30 hr to 36hr (Process III) and then increased to 44 hr (Process IV). This time dependency of *c*-values is similar to that of Stream 1.

#### Stream 3

The *s*-value was not constant but oscillated between 14.5S (around 38 hr) and 16.3S (around 26 hr). In addition, the *c*-values also oscillated with 24 hrs period: The *c*-values became minimum around 28 hr and maximum around 38hr, showing that the *s*and *c*-values oscillate with opposite phases.

To confirm the existence of these streams, we also performed time-resolution size exclusion chromatography (tr-SEC), which focused on the elution time covering Streams 2 and 3 (see detail in Material and Methods). As shown in Fig.S3, the tr-SEC also shows the streams corresponding to Streams 2 and 3. The streams measured by AUC have been confirmed. As the next step, it is crucially necessary to identify the components corresponding to the streams and found a new picture consistent with the AUC result.

### Assignment of components of streams

To assign the components in the streams with their *s*-values, we performed six AUC experiments, four single solutions of KaiA (A_2_ oligomer), KaiB (B_4_ oligomer), KaiC (C_6_ oligomer) and B_6_C_6_ complex, and for two titrations of KaiA+KaiC (A_2_+C_6_) and KaiA+B_6_C_6_ complex (A_2_+B_6_C_6_). The AUC profiles are shown in Figs.S4–S6. These experiments revealed the contributors of three streams as follows:

#### Stream 1

The *s*-value of Stream 1 was 2.83 ± 0.13S (Fig.3(a)). As shown in Fig.S4, the *s*-values of A_2_ and B_4_ oligomers are 3.563 ± 0.003S and 2.927 ± 0.004S, respectively. Therefore, Stream I is assigned to be B_4_ oligomers.

#### Stream 2

The *s*-value of Stream 2 is 11.75 ± 0.08S but there are no homo-oligomers nor complexes with that *s*-value (Fig.S4). Then, we focused on the titration experiment of KaiA and KaiC. As shown in Fig.S5(a), by adding KaiA (A_2_ oligomer) to C_6_ oligomer, the peak of C_6_ oligomer at 11.2S shifted to the higher *s*-value and the peak position reaches around 12.4S, which is supposed to be the *s*-value of A_2_C_6_ complex. Therefore, the peak between 11.4S and 12.4S corresponds to the dissociation-association equilibrium state between C_6_ and A_2_C_6_ (Fig.S5(b)). The molar ratio of C_6_ and A_2_C_6_ in Stream 2 is found to be [C_6_]:[A_2_C_6_] = 0. 47: 0.53, following the procedure proposed by Schuck et al [20].

#### Stream 3

The *s*-value of Stream 3 was not static but oscillated between 14.5S and 16.3S (Fig.3(a)) indicating that Stream 3 oscillated between the complexes larger than B_6_C_6_ and A_2_C_6_ complexes (*s* = 12.4S, Figs.S3 and S4) at least. Namely, Stream 3 corresponds to the A*_n_*B_6_C_6_ complexes with the dynamically oscillating association number of KaiA (*n*-value). To elucidate the *n*-values, we focused on the KaiA+B_6_C_6_ titration experiment. As shown in Fig.S6(a), by adding KaiA to B_6_C_6_, the peak of B_6_C_6_ oligomer at 12.4S shifted to the higher *s*-value pointed by the colored arrows, indicating that A*_n_*B_6_C_6_ complexes were generated with the larger *n*-value. Figure S6(b) shows the molecular weight as a function of *s*_20,*w*_ (12≤ *s*_20,*w*_ ≤18). Here the molecular weights of the peaks provided with SEDFIT software [21] are expressed with the colored closed circles and black curve shows the result of least squares fitting with a quadratic function. The molecular weights of A*_n_*B_6_C_6_ complexes (*n*=2, 4, 6, 8, 10, 12) are also shown with broken lines. From the intersections between the black solid and broken lines, we can know the *s*-values of the A*_n_*B_6_C_6_ complexes: The molecular weights and the *s*-values of the A*_n_*B_6_C_6_ complexes are listed in Table 1. The oscillation range of Stream 3 (14.5S and 16.3S) highlighted by the shadow zone revealed that Stream 3 mainly corresponds to the oscillation between *n*=4 (A_4_B_6_C_6_) and *n*=10 (A_10_B_6_C_6_). Figure 4(a) shows the relation between the Streams 2 and 3 and the assigned *s*-values of oligomers/complexes.

**Figure 4.**
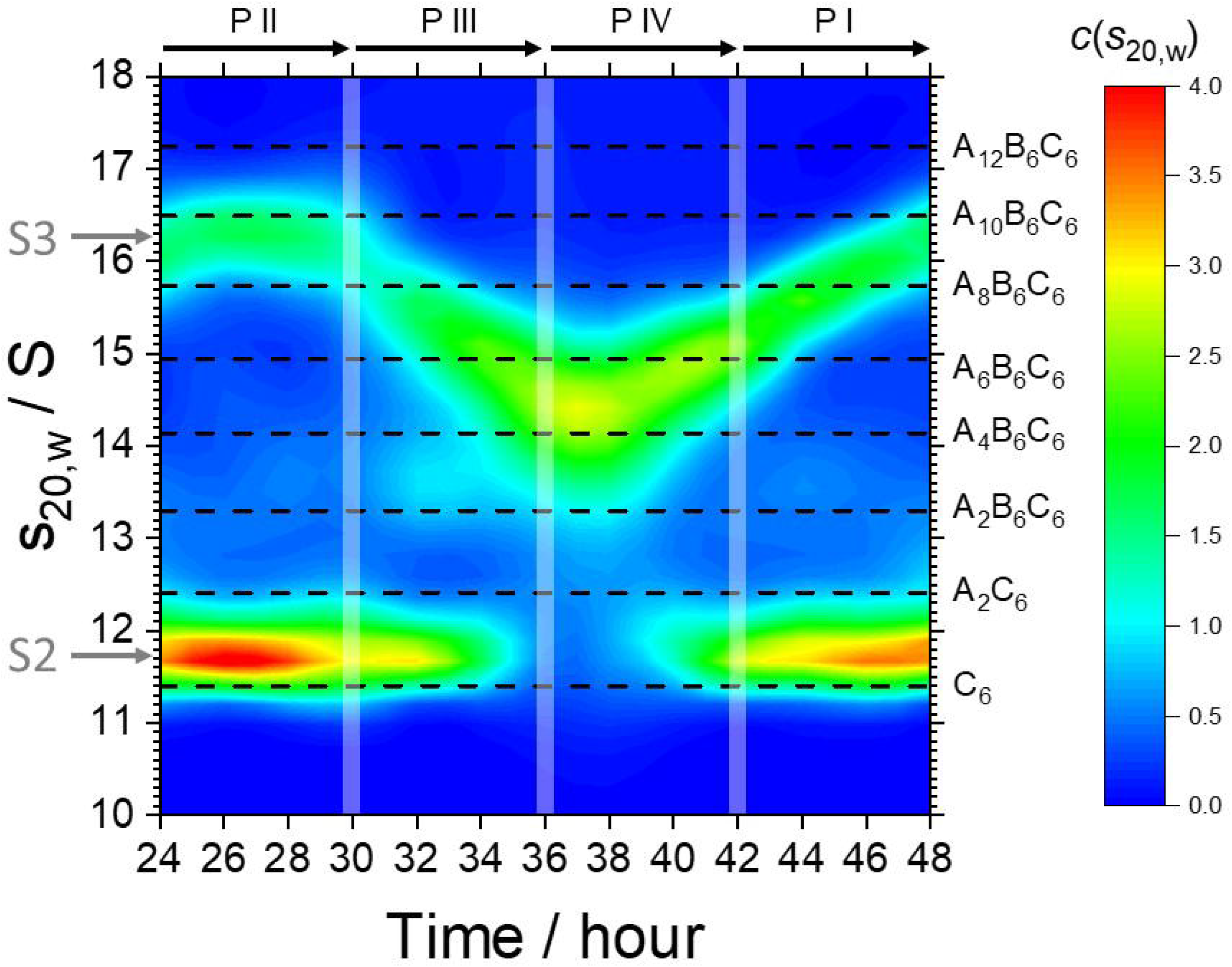
Relation between two streams and the formed oligomer/complexes on twodimensional map of time evolution of AUC-spectra. Streams are indicated by grey arrows at the beginning points and *s*-value (*s*_20,*w*_) corresponding to the oligomer/complexes are also drawn with black broken lines. See main text about the relation between *s*-value (*s*_20,*w*_) and the oligomer/complexes.

**Table 1.** Characteristic parameters of all components in Kai-clock system.

| Oligomer<br>/Complex | B<sub>4</sub> | A<sub>2</sub> | C<sub>6</sub> | A<sub>2</sub>C<sub>6</sub> | B<sub>6</sub>C<sub>6</sub> | A<sub>x</sub>B<sub>6</sub>C<sub>6</sub> |  |  |  |  |  |
| --- | --- | --- | --- | --- | --- | --- | --- | --- | --- | --- | --- |
|  |  |  |  |  |  | 2 | 4 | 6 | 8 | 10 | 12 |
| Mw | 46 | 65 | 348 | 413 | 417 | 481 | 547 | 612 | 678 | 743 | 808 |
| $S_{20,w}$ | 2.93 | 3.56 | 11.2 | 12.3 | 12.4 | 13.3 | 14.1 | 14.9 | 15.7 | 16.5 | 17.2 |
| $S_{20,w,min}$ | 1.18 | - | 10.1 | - | - | 12.8 | 13.7 | 14.5 | 15.4 | 16.1 | 16.9 |
| $S_{20,w,max}$ | 3.50 | - | - | 12.8 | - | 13.7 | 14.5 | 15.4 | 16.1 | 16.9 | 17.6 |

### Kinetics of the components in the oscillation system

To clarify the kinetics of components, B_4_, A_2_, C_6_, A_2_C_6_, and A_n_B_6_C_6_ (*n*=2,4,6,8,10,12), we estimated time evolutions of their numbers based on the *s*-, *c*-values and their molecular weights. The number of the x-components, *N*_x_, was estimated with *N*_x_ = ∑*_s_c*(*s*) /*m*_x_. Here, the sum was taken over the *s*-range corresponding to the *x*-components, and *m_x_* was the molecular weight: The corresponding *s*-range (*s*_20,*w*,min_ < *s*_20,*w*_ < *s*_20,*w*,max_) and *m_x_* are listed in Table 1. Figure 5 shows the time evolutions of number distributions of all components. This figure shows the kinetics of all components in a working Kai-clock system. However, because these number distributions were provided by only AUC data, we should examine the distributions with different methods. As the first examination, we checked the consistency of the number distributions with the SAXS result. For this purpose, we noticed that the *I*(0) is roughly proportional to a sum of the square of molecular weight of all components in the system, 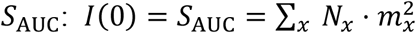. we calculated the time evolution of S_AUC_ utilizing {*N_x_*} provided by the AUC experiments (Fig.5). Figure 6 compared it with the time evolution of the experimental *I*(0). The time evolution of S_AUC_ well reproduced the that of the experimental *I*(0), meaning that the number distribution provided by AUC correctly expresses the real distribution in the Kai-clock system.

**Figure 5.**
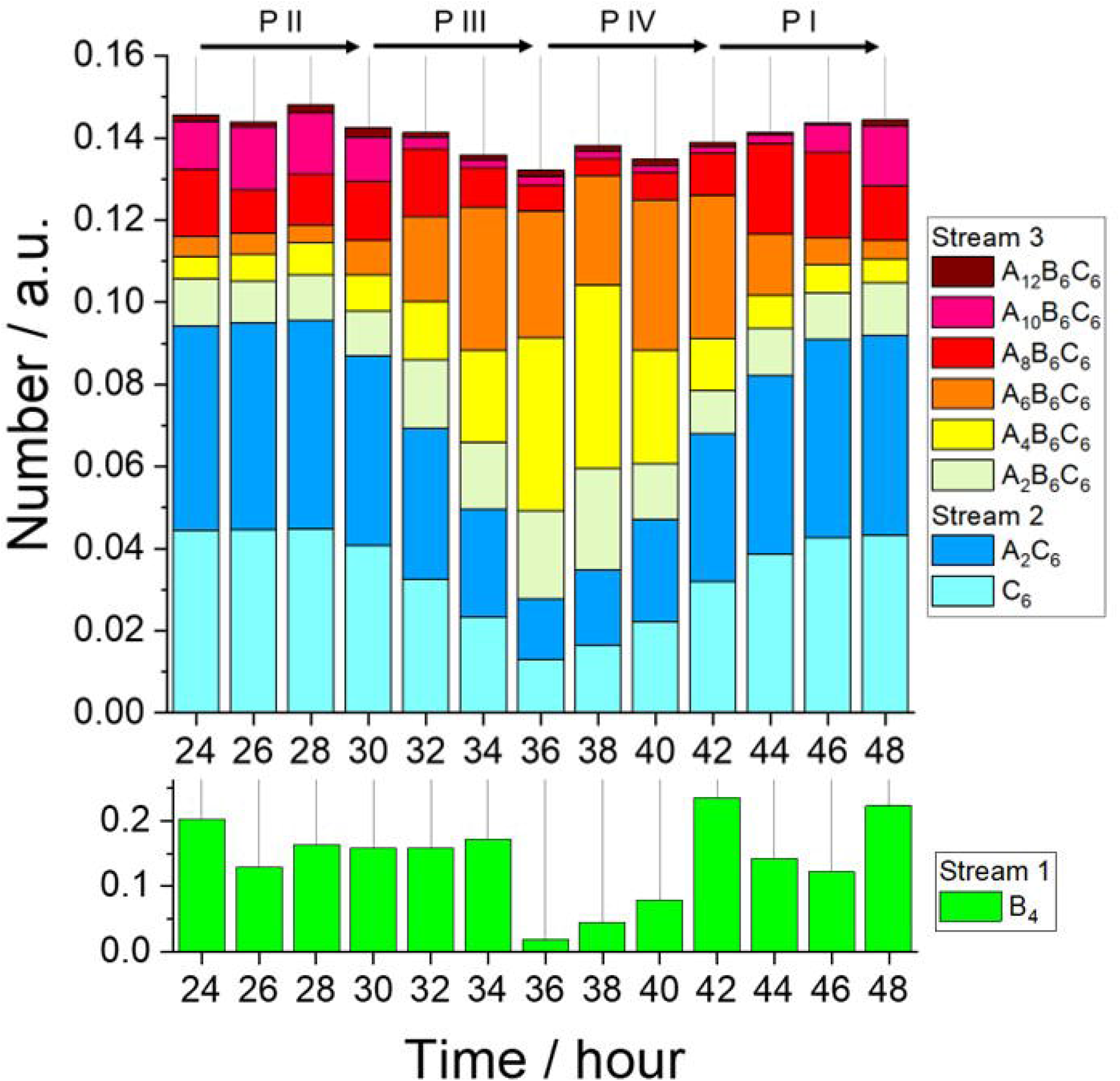
Time evolution of the number distributions of all components during a working Kai-clock. The number of each component is expressed with the length of bar: The scales of number are different between the lower (Stream 1) and upper (Streams 2 and 3) panels.

**Figure 6.**
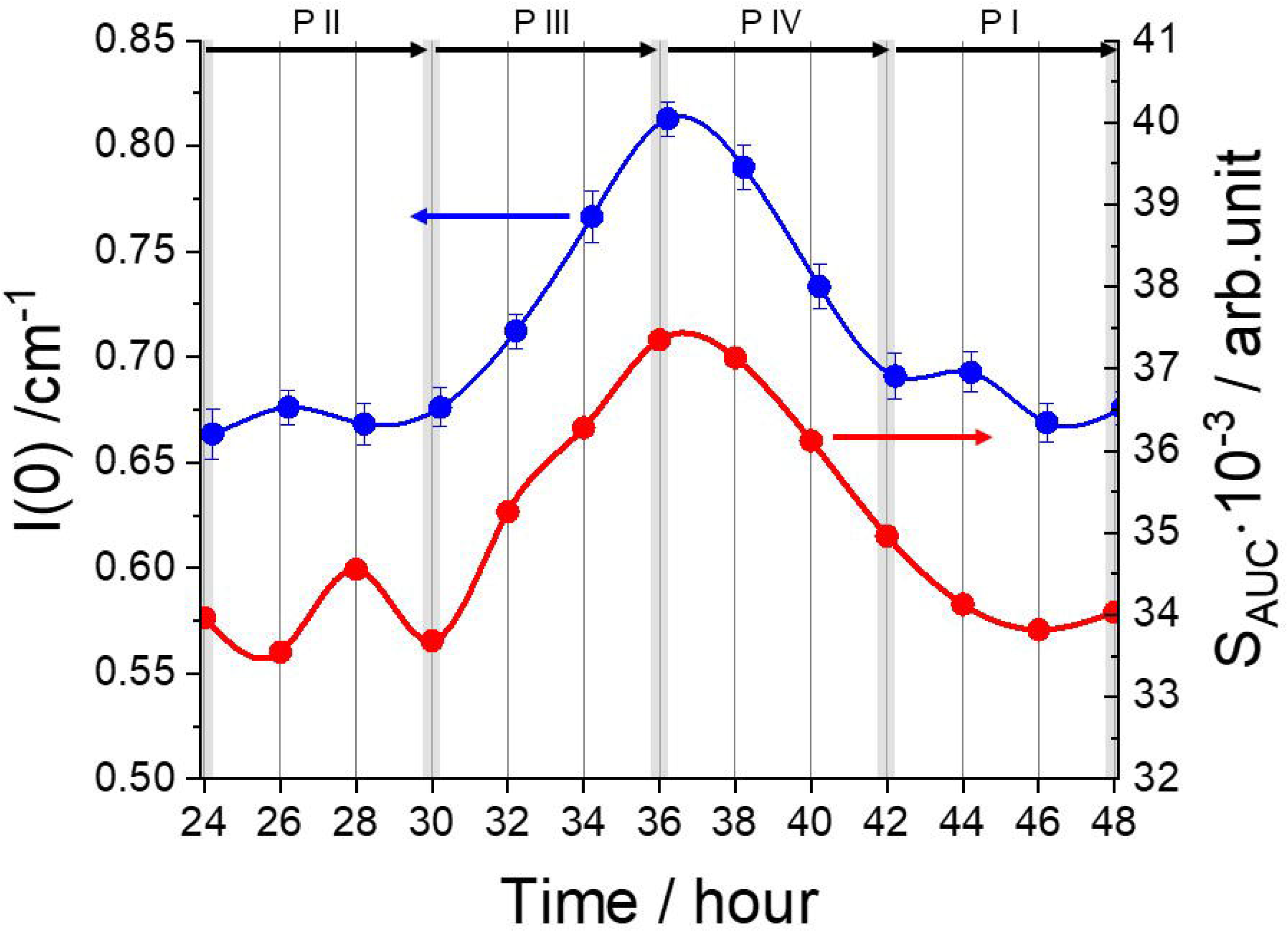
Relation between the measure forward SAXS intensity and S_AUC_ calculated with the AUC profiles. S_AUC_ (red circles) is expected to be proportional to the forward SAXS intensity (blue circles). Lines are eye guides.

As the second examination, we focused on kinetics of each molecule, KaiA molecule, KaiB molecule and C_6_ unit (KaiC). Figure S7(a)-(c) shows the numbers in the streams and their total numbers of all molecules/unit in a working Kai-clock. Even though the AUC experiments do not have the enough resolution for analyzing the precise numbers of small oligomers, such as B_4_ and A_2_, the total numbers for all molecules were almost hold. The averaged total numbers of KaiA, KaiB and KaiC are 0.46 ± 0.03, 0.98 ± 0.20 and 0.84 ± 0.03, of which ratios, [KaiA]/[KaiC] = 0.54 ± 0.04 and [KaiB]/[KaiC] = 1.16 ± 0.24, are almost agreements with the initial charged ratios, [KaiA]/[KaiC] = 0.46 and [KaiB]/[KaiC] = 0.13. Accordingly, AUC properly detected the kinetics of all oligomers in a working Kai-clock. Furthermore, we should emphasize two points about kinetics of C_6_ oligomers as clearly shown in Fig.S7(c): one is that the kinetics of C_6_ oligomer is oscillation between Streams 2 and 3 and the other is that the total number was hold. This means that the C_6_ oligomers are not decomposed and working as a base unit for a Kai-clock system.

As the final examination, we checked the consistency of kinetics of KaiA analyzed by the AUC result with tr-CM-SANS experiment (see the detailed theory of CM-SANS in Supporting note 2). As shown in Fig. 5, there are the largest A_n_B_6_C_6_ complexes (*n*→8~10) in Processes I and II whereas the smallest A_n_B_6_C_6_ complexes (*n*→2~4) in Processes III and IV. Here, we focused on kinetics of KaiA. In Processes I and II, there are small number of A_n_ oligomers (*n*→8~10) in Stream 3 and the large number of A_2_ oligomers in Stream 2 whereas in Processes III and IV, there are the large number of A_n_ oligomers (*n→*2~4) in Stream 3 and small number of A_2_ oligomers in Stream 2. We checked the consistency of the above-mentioned distributions of KaiA with the CM-SANS: Because the CM-SANS can selectively observe KaiA protomers in ABC complex and A_2_ oligomers in a working Kai-clock (Fig.S10), we can examine the KaiA distributions in a working Kai-clock (see the detailed method of CM-SANS in Supporting note 2). With the same manner of the first examination, we calculated the time evolution of 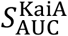 utilizing the above-mentioned protomers in ABC complex and A_2_ oligomers: 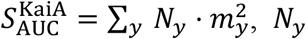 and *m_y_* are *y*-mer (y=2,4,6,8,10,12) and its molecular weight, respectively. Interestingly, as shown with red circles in Fig.7, the calculated 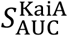 oscillated with the phase opposite to those of S_AUC_ and the experimental SAXS 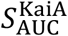 becomes maximum around Processes I and II and minimum around Processes III and IV. As shown with blue circles in Fig.7, the time evolution of *I*(0) of the CM-SANS experiment well reproduced that of the calculated 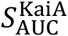. Therefore, the kinetics of KaiA reveled with the AUC results is also confirmed with the CM-SANS experiment.

**Figure 7.**
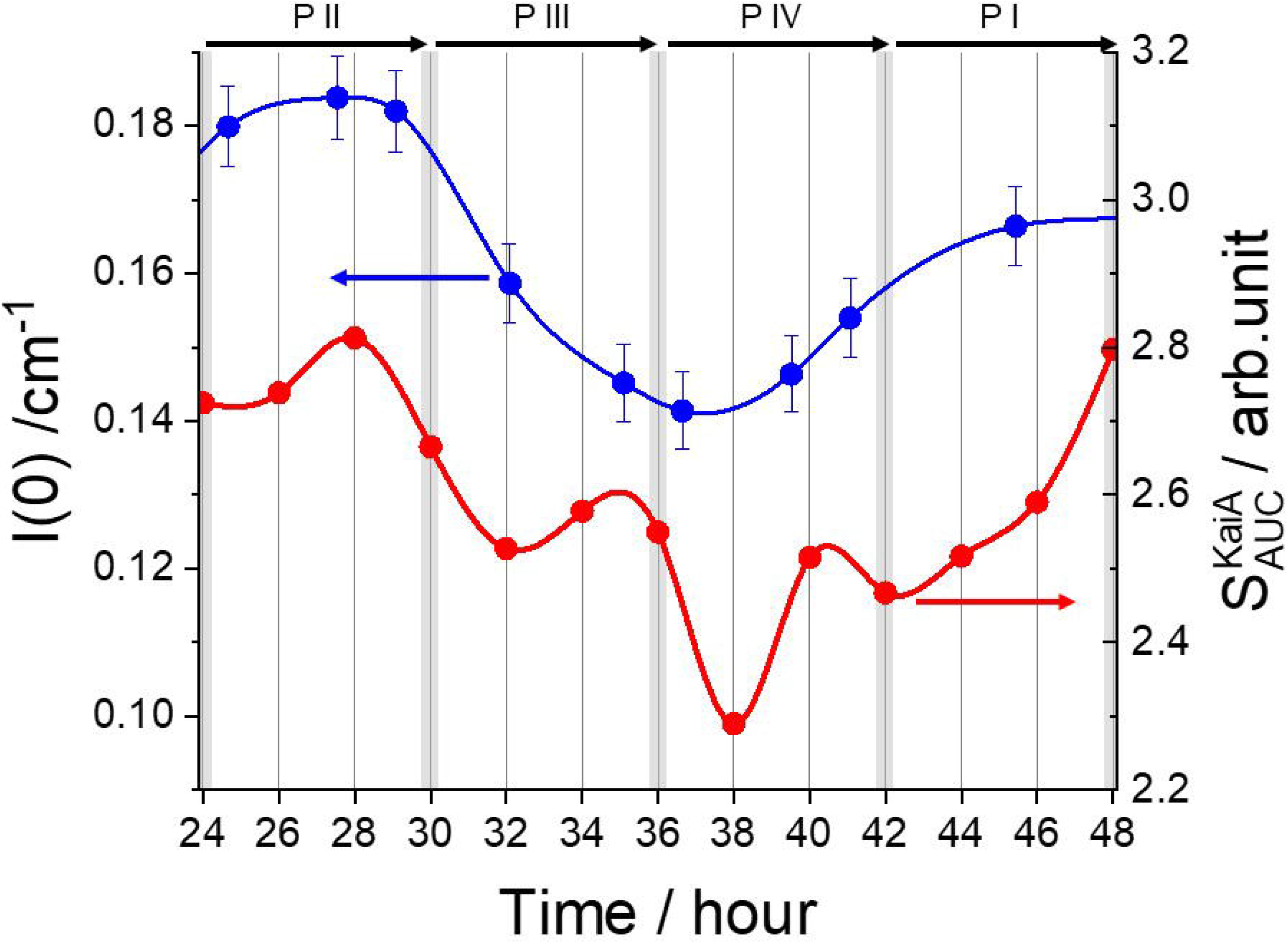
Relation between the measure forward CM-SANS intensity and 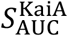 calculated with the AUC profiles. 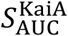 (red circles) is expected to be proportional to the forward SAXS intensity (blue circles). Lines are eye guides.

## DISCUSSION

Our results disclosed that Kai-clock has not a single line time oscillation, as shown in Fig.S1, but is composed with corelating three oscillation streams: Stream 1 is the number oscillation of B_4_ oligomer, Stream 2 is the oscillation of total number of components in the equilibrium state between C_6_ and A_2_C_6_, and Stream 3 is the coupled oscillations with total number and association number of KaiA of ABC complex.

Taking consideration of the relation with three streams, we summarize the time transition of components in the Kai-clock oscillation in the Fig.8. To explain the transition, we select four time points, Points D, A, B and C where *I*(0) exhibits maximum, the PosD does minimum, *I*(0) does minimum, the PosD does maximum. In addition, the box size and the position on y-axis approximately exhibit the number of components and the average *s*-values in the streams, respectively.

**Figure 8.**
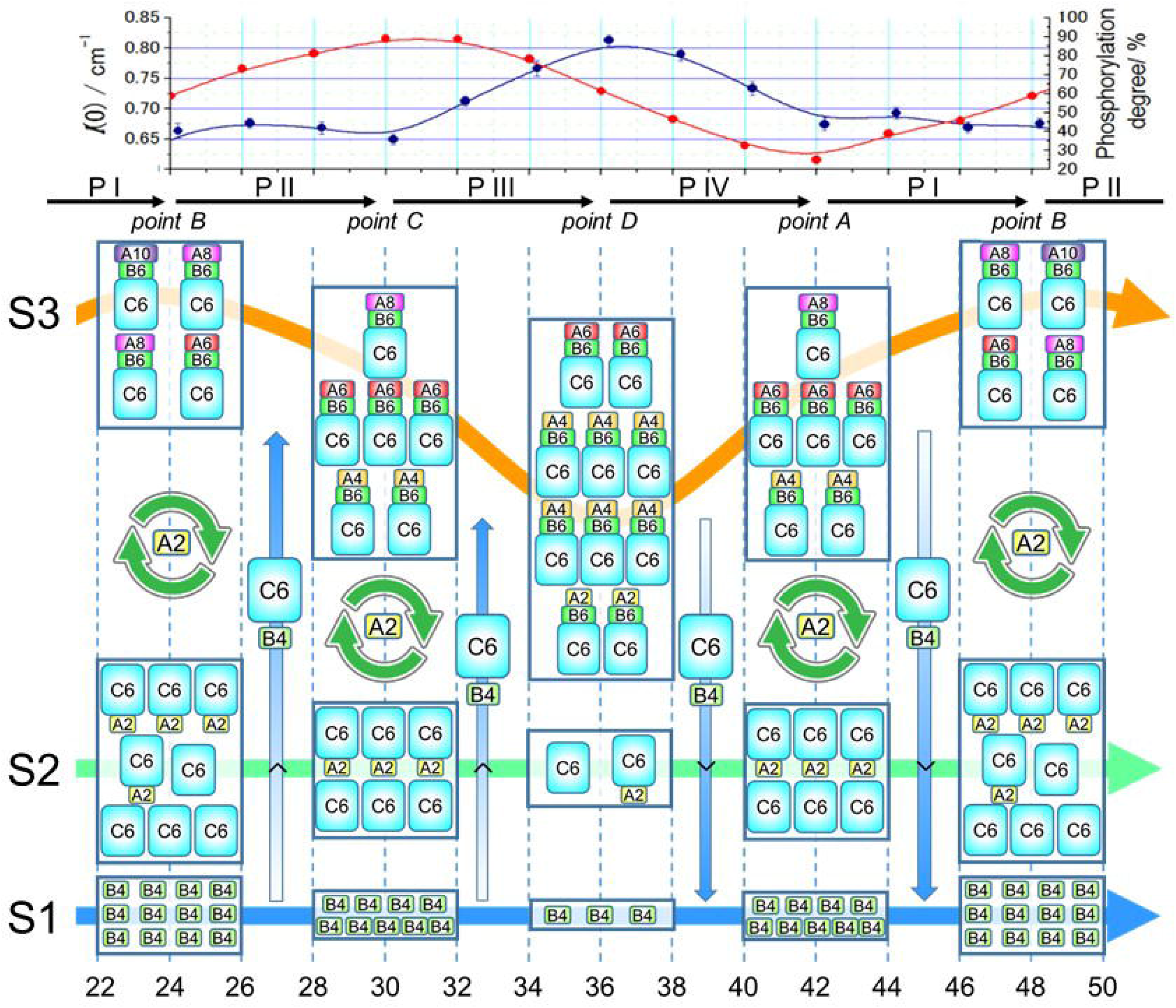
Schematic view of the kinetics of all components in a working Kai-clock system. In the center panel, x- and y-axes correspond to elapsed time and *s*-value in AUC profile. Streams 1, 2, and 3 are expressed with thick blue, green, and orange lines, respectively. The representative components in streams are drawn in the boxes on the streams at points A-D. The positions and sizes of boxes roughly express the averaged *s*-values and numbers of constituents, respectively. The combinational marks with rotational arrows and A_2_ exhibit the existence of the distribution equilibrium of A_2_ oligomers between Streams 2 and 3. The thick white-blue arrows with C_6_ and B_4_ also denote the moving directions of B_4_ and C_6_ oligomers in the corresponding processes shown with the solid black arrows. The upper panel shows the simultaneous oscillations of the forward SAXS intensity and phosphorylation degree of KaiC.

### Point D

In Stream 3, the largest number of ABC complexes are formed. The average association number of KaiA in ABC complex is minimum (~4) since the number of KaiA providing for ABC complexes is limited. In Stream 2, the numbers of the components, C_6_ and A_2_C_6_, become minimum, which suppresses the phosphorylation of KaiC and makes the speed of dephosphorylation of KaiC largest.

### Process IV

In Stream 3, KaiC protomers in ABC complexes are dephosphorylated, accompanying with the decomposition of ABC complexes as well. Therefore, the A_2_, B_4_ and C_6_ oligomers are released and the numbers of the components belonging to Streams 1 and 2 are increasing whereas that of Stream 3 is decreasing. Even though C_6_ units are being transferred from Stream 2 to Stream 3 by formation of ABC complexes, named as UpC_6_, the amount is smaller than that being transferred from Stream 3 to Stream 2 by decomposition of ABC complexes, named as DownC_6_.

### Point A

In Stream 2, the dephosphorylated C_6_ oligomers make A_2_C_6_ oligomers associating with the A_2_ oligomers, meaning that the phosphorylation of KaiC is activated. The amount of A_2_ released by decomposition of ABC complex in Process IV is much larger than those used for the formation of A_2_C_6_. As a result, the released A_2_ returns to Stream 3 and associates to ABC complex, inducing the increase of the average association number of KaiA in ABC complex (4→6). This also means that there is an equilibrium in KaiA between Streams 2 and 3.

### Process I

In Stream 3, the dephosphorylation of KaiC protomers continues, making the decomposition of the ABC complexes. In Stream 2, A_2_C_6_ phosphorylates C_6_ protomer and then the phosphorylated C_6_ oligomer forms an ABC complex via the formation of B_6_C_6_ complex. In this process, the amount of UpC_6_ is still smaller than that of Down C_6_. As a result, the numbers of the components belonging to Streams 1 and 2 are still increasing whereas that of Stream 3 is decreasing.

### Point B

In Stream 2, the largest number of C_6_ units (homooligomers and protomers in A_2_C_6_ oligomers) belongs and, therefore, the phosphorylation of KaiC is mostly activated. With the same reason described in Point A, the number of ABC complexes is decreased but the average association number of KaiA in ABC complex is increased (6→8~10).

### Process II

In Stream 2, A_2_C_6_ produces the phosphorylated Kai C molecules in C_6_ protomer, subsequently they form ABC complexes with B_4_ in Stream 1 as well. Contrary to Process I, in this process, the amount of DownC_6_ is superior to that of UpC_6_. As a result, the numbers of the components belonging to Streams 1 and 2 are decreasing whereas that of Stream 3 is increasing.

### Point C

The increased number of ABC complex is same as the decreased number of C_6_ units (homooligomers and protomers in A_2_C_6_ oligomers). Considering the amount of providing A_2_ from Stream 2 to Stream 3, total number of KaiA in ABC complex is increased but the average association number is decreased (8~10→6).

### Process III

In Stream 2, A_2_C_6_ continuously produces the phosphorylated Kai C molecules in C_6_, subsequently the phosphorylated C_6_ oligomers form ABC complexes with B_4_ in Stream 1 as well. Same as Process II, in this process, the amount of UpC_6_ is superior to that of DownC_6_. As a result, the numbers of the components belonging to Streams 1 and 2 are decreasing whereas that of Stream 3 is increasing. Then, the Kai-Clock system returns back to Point D.

We noticed that the ratio of C_6_ and A_2_C_6_ in Stream 2 holds 1:1. This is supported by the evidence that the peak of Stream 2 was not changed. As the mechanics of this hold, the surplus KaiA dimers are absorbed by Stream 3 through the formation of ABC complex. Otherwise, the free KaiA dimers associate with C_6_ in Stream 2 and produce more A_2_C_6_. In other words, this management KaiA works to make the yield of the phosphorylated KaiC in Stream 2 constant. This could be important for stable oscillation.

## ACKNOWLEDGEMENTS

This work was supported by MEXT/JSPS KAKENHI Grant Numbers (JP18H05229 and JP18H03681 to M.S. JP19K16088 and JP21K15051 to K.M., JP17K07361, JP19KK0071 and JP20K06579 to R.I., JP17K07816 to N.S.) and by Research Fund for Young Scientists in Kyoto University and Fund for Project Research in Institute for Integrated Radiation and Nuclear Science, Kyoto University (KURNS) to K.M. This work was also partially supported by the project for Construction of the basis for the advanced materials science and analytical study by the innovative use of quantum beam and nuclear sciences in KURNS. The study was partially supported by the Platform Project for Supporting Drug Discovery and Life Science Research (Basis for Supporting Innovative Drug Discovery and Life Science Research (BINDS)) from AMED.

## AUTHOR CONTRIBUTIONS

M.S., H.Y. and K.K. designed research. M.S., K.M., Y.Y., R.I. and N.S. performed simultaneous measurements of time-resolved AUC, SAXS and SDS-PAGE. M.S., K.M. and R.I. also performed simultaneous measurements of time-resolved AUC and CM-SANS. K.M. and Y.Y. performed titration measurements with AUC. K.M., Y.Y. and H.Y. prepared for samples. All authors wrote the paper.

## DATA AVAILABILITY

The datasets generated and analyzed during the current study are available from the corresponding authors on reasonable request.

## MATERIALS and METHODS

### Expression and purification of Kai proteins

KaiA, KaiB, and KaiC from *Synechococcus sp*. PCC 7942 were expressed in *Escherichia coli*. KaiA was generated as hexahistidine (his)-tagged recombinant protein and purified after the cleavage of the his-tag as described previously [22]. KaiB was generated as a glutathione S-transferase (GST)-tagged recombinant protein and purified after the cleavage of the GST-tag as described previously [23]. KaiC was generated as a Strep-tagged recombinant protein and purified as described previously [24].

For the preparation of the deuterated proteins, the bacterial cells were grown in M9 minimal D_2_O media containing deuterated glucose (1,2,3,4,5,6,6-D7, 98%, Cambridge Isotope Laboratories, Inc.) according to the previous study [25].

All proteins were finally purified by size exclusion chromatography (SEC) with an eluent containing 50 mM sodium phosphate (pH7.8), 150 mM sodium chloride, 5mM magnesium chloride, 0.5 mM EDTA, 1 mM dithiothreitol, 3 mM adenosine triphosphate (ATP), 50 mM glutamic acid, and 50 mM arginine.

### Time-resolved and simultaneous AUC-SAXS-phosPAGE

A reservoir solution was made by a mixture of KaiA, KaiB, and KaiC: the partial concentrations of KaiA, KaiB, and KaiC were 0.45 mg/mL, 0.45 mg/mL, and 1.8 mg/mL, respectively. Every 2 hr right after the mixture until 60hr, SAXS and contrast matching (CM)-SANS were measured for 36 and 90 min, respectively. At the same time, the reduction treatment was subjected to the sample for sodium dodecyl sulfate-polyacrylamide gel electrophoresis (SDS-PAGE). After 24 hr from the mixture, AUC and SEC were measured for 80 and 30 mins, respectively, every 2hr in 24 hrs. The schematic image of time course of AUC, SAXS, CM-SANS, SEC, and SDS-PAGE was shown in Fig.S8.

### AUC measurement

Sedimentation velocity-AUC measurements were performed with ProteomeLab XL-I (Beckman Coulter Inc., Brea, CA, USA). The optical path and the volume of the cell were 12 mm and 400 μL, respectively. All measurements were performed using Rayleigh interference optics at 60,000 rpm. The temperature was kept at 30°C for all measurements. The AUC profile, weight concentration distribution *c*(*s*_20,*w*_), was obtained by fitting the time evolution sedimentation data with Lamm formula using SEDFIT software (version 15.01c) (http://www.analyticalultracentrifugation.com/sedfit.htm) [21]. The sedimentation coefficient was normalized to be the value at 20 °C in pure water, *s*_20,w_. The molecular weight was calculated using the *s*_20,w_-value and the friction ratio *f/f_0_* [^26^ref x7].

### SAXS measurement

SAXS measurement was performed with NANOPIX (RIGAKU Co., Ltd., Tokyo, Japan) equipping with the point-focused generator of a Cu-Kα source (wavelength = 1.54 Å) and HyPix-6000 detector. The sample-to-detector distance were set to 1330 mm and then the covered *q*-range was from 0.01 Å^-1^ to 0.2 Å^-1^. The observed SAXS intensity was corrected for background, empty cell and buffer scatterings, and transmission factors and subsequently with SAngler software (http://pfwww.kek.jp/saxs/SAngler.html) [27]. The unit of scattering intensity was converted to the absolute scale by referring to a standard scattering intensity of pure water at 20 °C (1.632 × 10^2^ cm^-1^) [28]. The forward scattering intensity *I*(0) was obtained with Guinier analysis.

### CM-SANS measurement

Contrast matching (CM)-SANS experiments were performed with QUOKKA instrument installed at the Australian Nuclear Science and Technology Organization (ANSTO), NSW, Australia. The SANS intensities were measured with the wavelength of 6.0 Å and sample-to-detector distance of 6.0 m: the covered *q*-ranges are 0.011 to 0.1 Å^-1^. The temperature was maintained at 30°C in the irradiation. The observed SANS intensity was corrected for background, empty cell and buffer scatterings, and transmission factors and subsequently converted to the absolute scale with the macro of IGOR Pro from NIST [^29^ref x10]. To observe the intensity from only KaiA, the CM-SANS experiment was conducted for the mixture of 100%-deuterated KaiA, hydrogenated KaiB, and hydrogenated KaiC in 40%-D_2_O (60%-H_2_O) buffer.

### SEC and SDS-PAGE analysis

SEC analysis was performed with Superose 6 increase 10/300GL column (GE Healthcare, Chicago, IL, USA). Flow rate was set at 0.75 mL/min. SDS-PAGE analysis was conducted according to the previous study [3]. The intensities of the bands were measured by densitometry using ImageJ software (National Institutes of Health, Bethesda, MD, USA).

### Titration experiments

To assign the components corresponding to Streams 2 and 3, two titration experiments were carried out.

#### (i) KaiA + KaiC

We used the dephosphorylation mimic KaiCAA in which phosphorylation sites S431 and T432 were substituted with alanine residues. AUC measurements were conducted for the KaiA + KaiC mixture solutions, in which the composition was [KaiA]: [KaiC] = *x*: 6 (*x* = 0 - 8). Here, the partial concentration of KaiC was fixed at 1.0 mg/mL.

#### (ii) KaiA + B_6_C_6_ complex

We used the phosphorylation mimic KaiCDT in which a phosphorylation site S431 was substituted with an aspartate residue. B_6_C_6_ complex was prepared according to the previous work [13]. AUC measurements were conducted for the KaiA + B_6_C_6_ mixture solutions, in which the composition was [KaiA]: [B_6_C_6_] = *x*: 6 (*x* = 0 - 18). Here, the partial concentration of B_6_C_6_ was fixed at 0.6 mg/mL.

