## Supplementary Information for "Orchestration of Proteins in cyanobacterial Circadian Clock System 1"

**Supplemental Figures**

**Supplemental Figure 1**

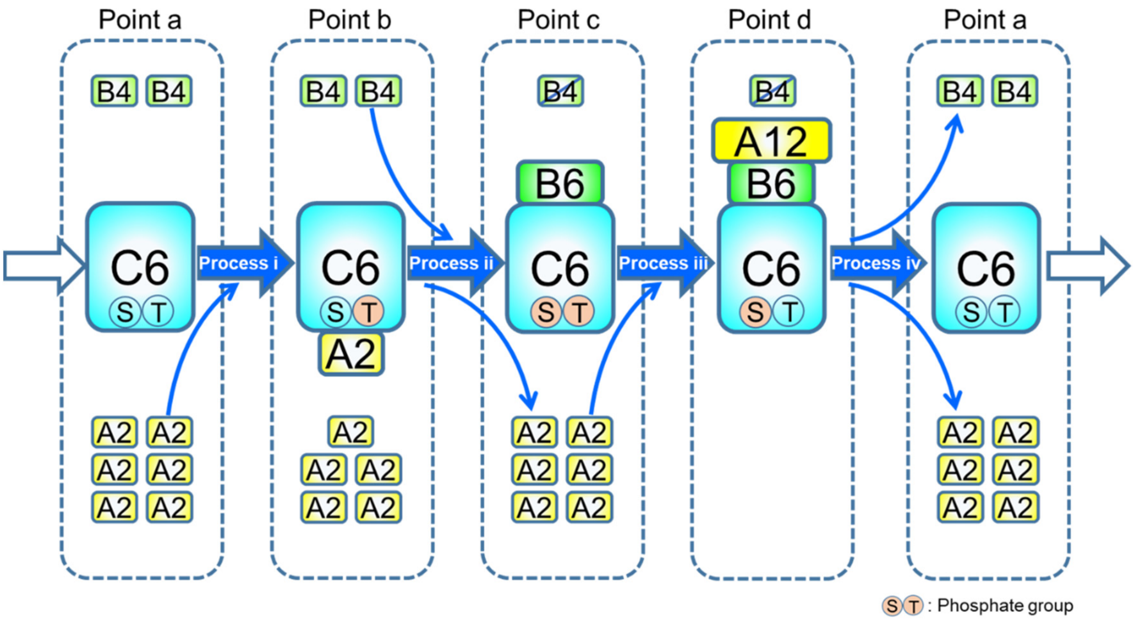


Figure S1. A proposed oscillation scheme in Kai clock system. Yellow, green, light blue squares show KaiA dimer/12-mer, KaiB tetramer/hexamer and KaiC hexamer, respectively: The numbers in squares express the association numbers in oligomer/protomer. Two phosphorylated cites, i.e. Ser431 and Thr432, in KaiC are shown with circles in a C6 square: The phosphorylation cycle is as follows: ST→pST→pSpT→pST→ST (p indicates a phosphorylated residue and colored with orange.) In dashed squares, representative components are drawn in at the timing points, Points a-d.

**Supplemental Figure 2**

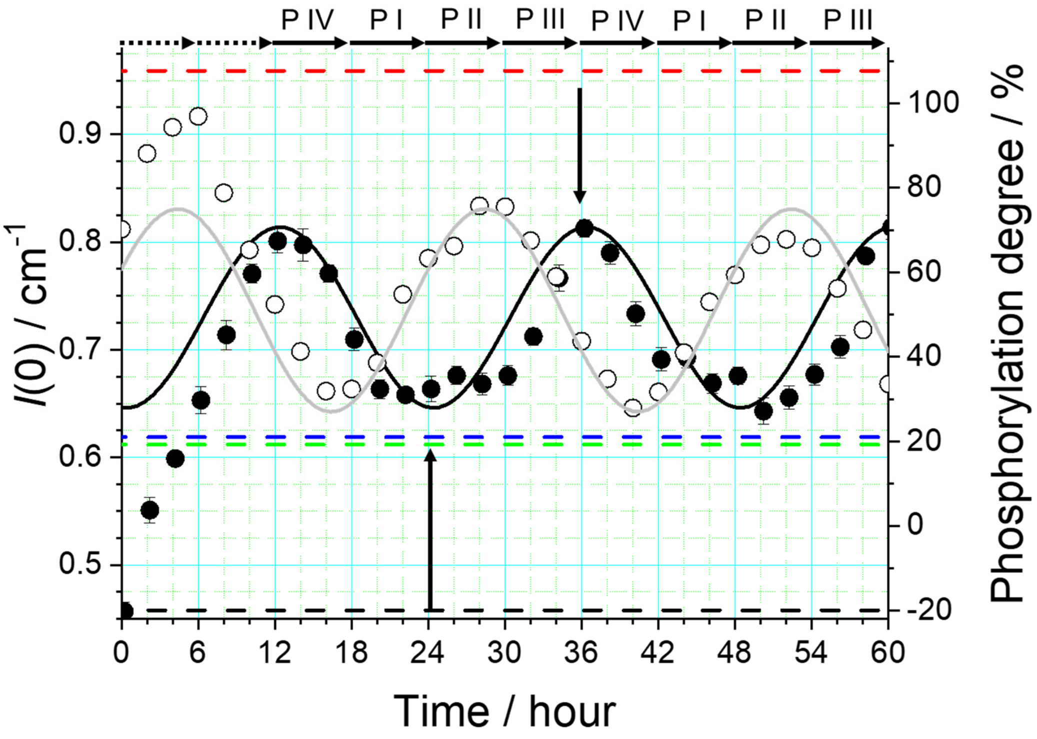


Figure S2. Time evolutions of the forward SAXS intensity and phosphorylation degree of KaiC during a working Kai-Clock. Closed and open circles show the forward SAXS intensity and phosphorylation degree of KaiC and black and grey lines are eye guides by trigonometric functions with a 24-hr period. Two vertical arrows point the minimum and maximum (right) forward SAXS intensities, $I_{\mathrm{min}}(0) \cong 0.65$ cm$^{-1}$ (left) and $I_{\mathrm{max}}(0) \cong 0.82$ cm$^{-1}$ (right), at 24 and 36 hrs, respectively. Black, blue, green and red broken lines express the forward SAXS intensities of comps 1-4, $I_1(0) = 0.458$ cm$^{-1}$, $I_2(0) = 0.619$ cm$^{-1}$, $I_3(0) = 0.612$ cm$^{-1}$, and $I_4(0) = 0.959$ cm$^{-1}$, respectively. (see main text in detail)

**Supplemental Figure 3**

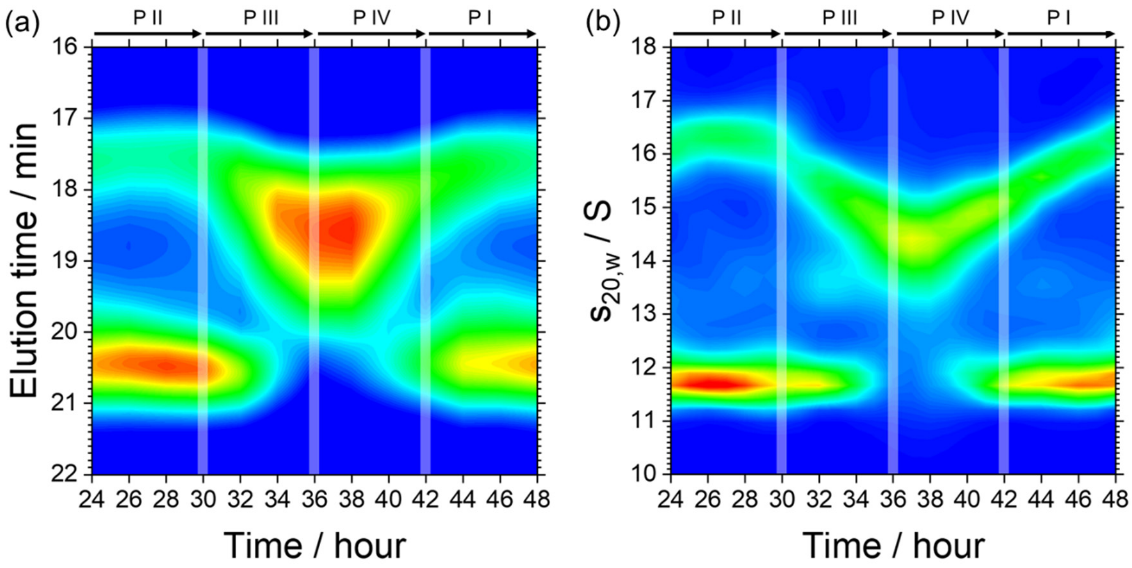


Figure S3. Time evolutions in a working Kai-clock: (a) Time-resolved size exclusion chromatography and (b) analytical ultracentrifuge (enlarged Fig. 2).

**Supplemental Figure 4**

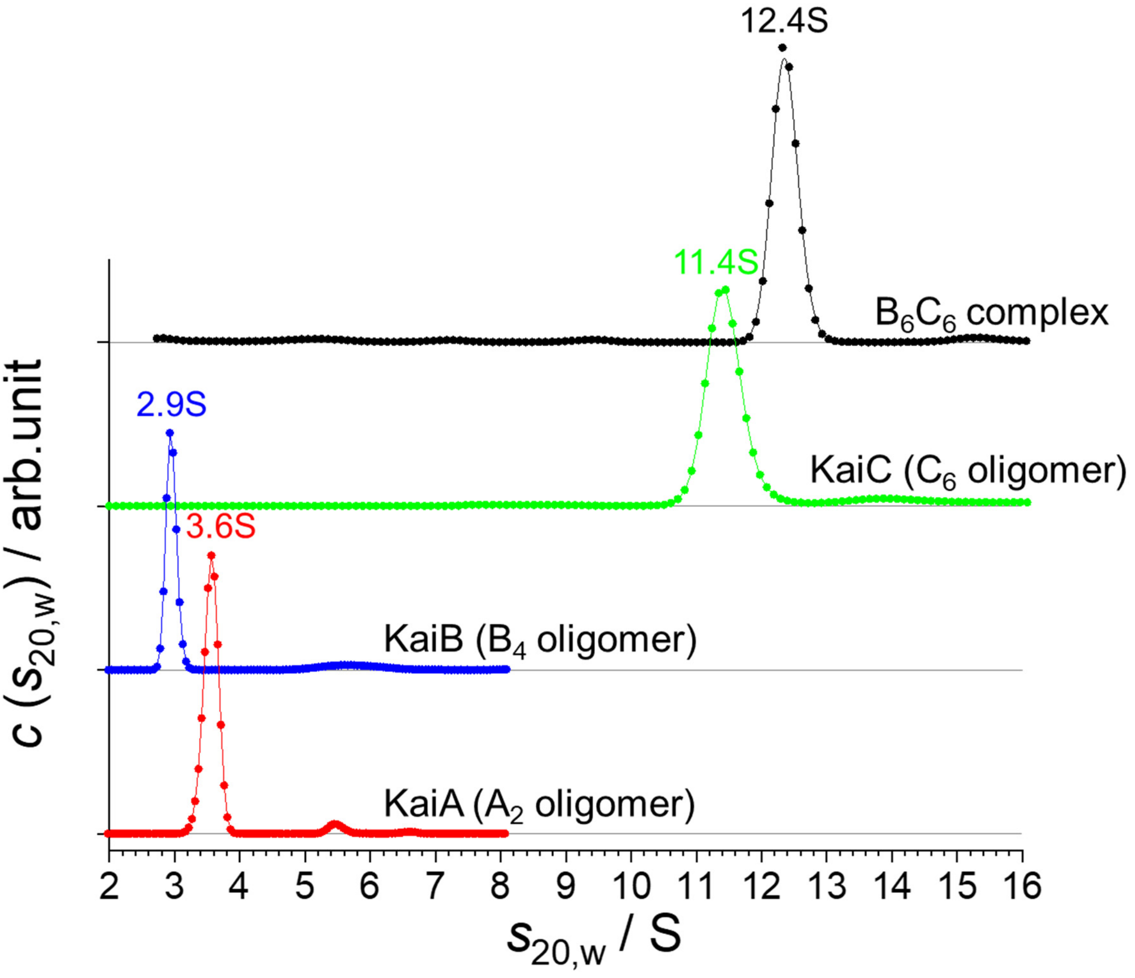


Figure S4. AUC profiles of solutions of three Kai proteins and $B_6C_6$ complex. (Red line) KaiA solution: a peak at 3.563 ± 0.003S corresponds to $A_2$ dimer, (Blue line) KaiB solution: a peak at 2.927 ± 0.004S corresponds to $B_4$ tetramer, (Green line) KaiC solution: a peak at 11.416 ± 0.003S corresponds to $C_6$ dimer, and (Black line) $B_6C_6$ complex solution: a peak at 12.357 ± 0.0013S corresponds to $B_6C_6$ complex.

**Supplemental Figure 5**

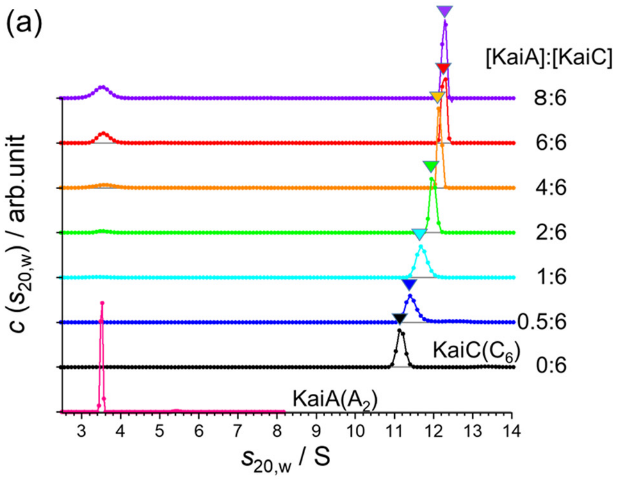


Figure S5. Titration of KaiA ($A_2$ dimer) and KaiC ($C_6$ hexamer) with AUC. (a) AUC profiles with different mixture ration between KaiA and KaiC, [KaiA] : [KaiC] = $x$ : 6 ($x$ = 0 - 8). Here, the partial concentration of KaiC was fixed at 1.0 mg/mL. The AUC profiles of KaiA and KaiC of solo-solutions are also shown as references. (b) The peak positions as a function of ratio of KaiA and KaiC. Black curve shows the result of least squares fitting with a Sigmoid function: The asymptotic value at the higher $s$ is 12.3S. The straight line and grey zone show the $s$-value of Stream 2 and the error, respectively.

**Supplemental Figure 6**

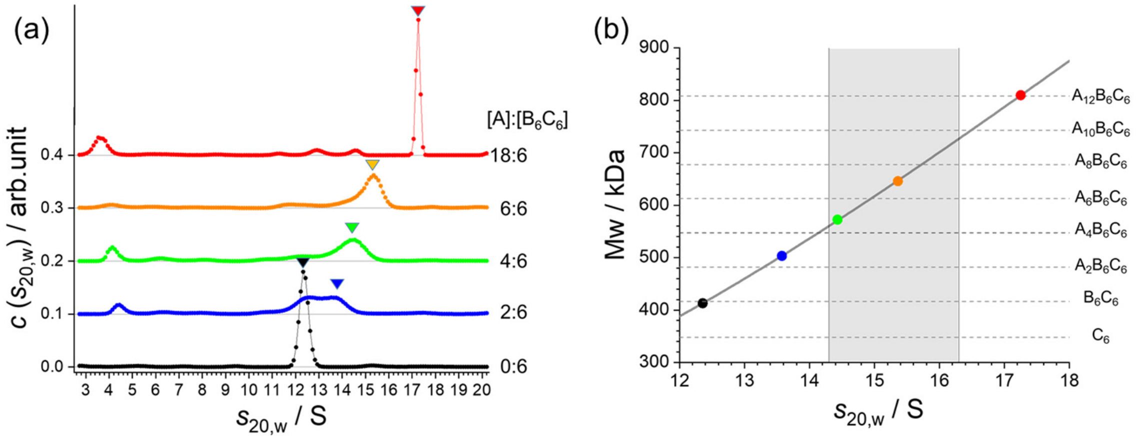


Figure S6. Titration of KaiA ($A_2$ dimer) and $B_6C_6$ complex with AUC. (a) AUC profiles with different mixture ration between KaiA and $B_6C_6$, [KaiA] : [$B_6C_6$] = *x* : 6 (*x* = 0 - 18). Here, the partial concentration of $B_6C_6$ complex was fixed at 0.6 mg/mL. The mutated $KaiC_{DT}$ in which a phosphorylation site S431 was substituted with an aspartate residue, are used as phosphorylation mimic KaiC. The AUC profiles of KaiA and KaiC of solo-solutions are also shown as references. (b) The molecular weights of ABC complexes as a function of *s* -value. Colored circles show the calculated molecular weights corresponding to the peaks in AUC profiles. Broken lines show the molecular weights of possible $A_xB_6C_6$ complexes (*x*=0, 2, 4, 6, 8, 10, 12) and $C_6$ hexamer. Black curve shows the result of least squares fitting with a quadratic function. The grey zone shows the *s*-value of Stream 3 including the error.

**Supplemental Figure 7**

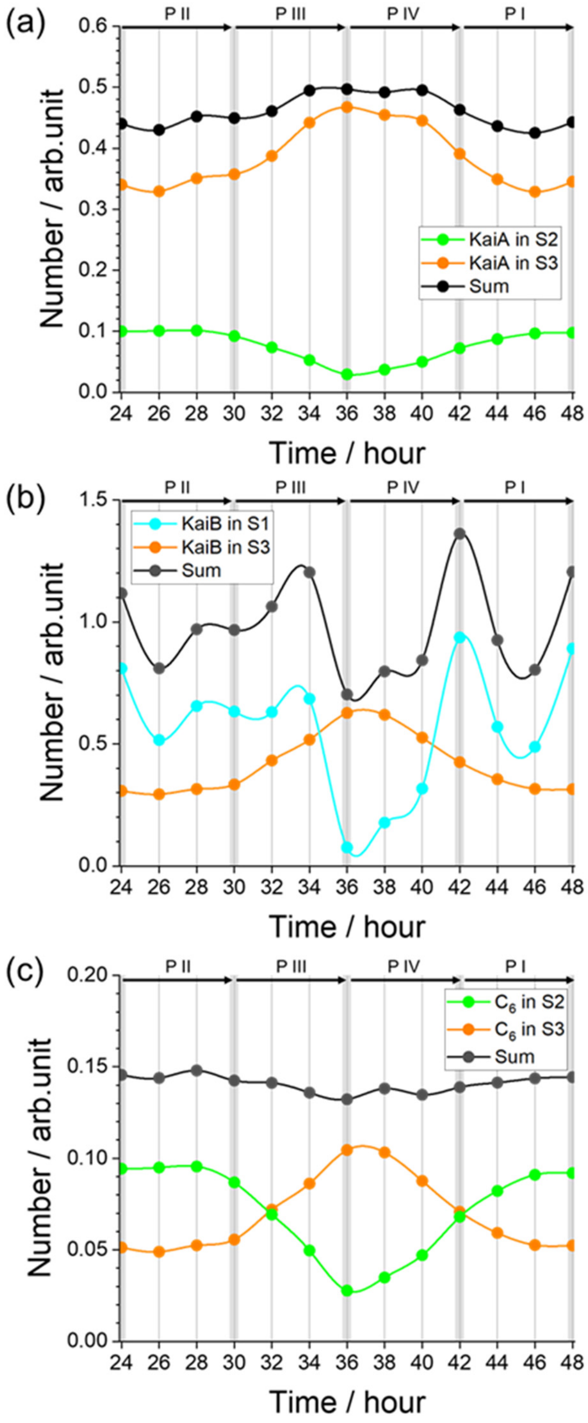


Figure S7. Time evolutions of the number distributions of KaiA, KaiB and $C_6$ hexamer in Streams. (a) KaiA distributions in Streams 2 and 3 and their sum. (b) KaiB distributions in Streams 1 and 3 and their sum. (C) $C_6$ hexamer distributions in Streams 2 and 3 and their sum.

**Supplemental Figure 8**

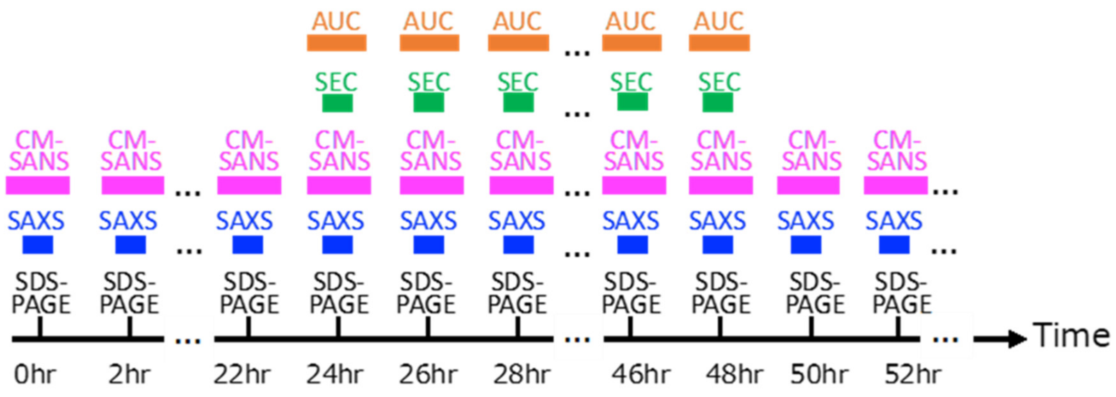


Figure S8. Time course of AUC, SEC, CM-SANS, SAXS, and SDS-PAGE (for measurement of phosphorylation degree) experiments.

**Supplemental note 1: Molecular ratio of $C_6$ oligomer and $A_2C_6$ complex**

As shown in Fig.S5, $C_6$ oligomer and $A_2C_6$ complex were observed as a merged peak in 11.2 S – 12.3 S depending on the mixing ratio of KaiA because the timescale of the association-dissociation ($A_2$+$C_6$ $\rightleftarrows$ $A_2C_6$) was much faster than that of AUC experiment. The sedimentation coefficient of the merged peak is represented as the weight-average of $C_6$ oligomer and $A_2C_6$ complex as follows [1].

$$s_{20,\mathrm{w}} = \frac{n_{\mathrm{C6}} M_{\mathrm{C6}} s_{20,\mathrm{w,C6}} + n_{\mathrm{A2C6}} M_{\mathrm{A2C6}} s_{20,\mathrm{w,A2C6}}}{n_{\mathrm{C6}} M_{\mathrm{C6}} + n_{\mathrm{A2C6}} M_{\mathrm{A2C6}}}\ , \qquad \text{(S1)}$$

where $n_{\mathrm{C6\ or\ A2C6}}$, $M_{\mathrm{C6\ or\ A2C6}}$, and $s_{20,\mathrm{w,C6\ or\ A2C6}}$ are molar ratios ($n_{\mathrm{C6}} = 1 - n_{\mathrm{A2C6}}$), molecular weights, and sedimentation coefficients of $C_6$ oligomer and $A_2C_6$ complex, respectively. From the saturating $s_{20,\mathrm{w}}$-value on the condition of excessive mixing ratio of KaiA, $s_{20,\mathrm{w,A2C6}}$ was estimated to be 12.3 S. Therefore, the molar ratio of $C_6$ and $A_2C_6$ in Stream 2 ($s_{20,\mathrm{w}}$ = 11.75 S) is given as $n_{\mathrm{C6}}$ = 0.47 and $n_{\mathrm{A2C6}}$ = 0.53, respectively.

**Supplemental note 2: Contrast Matching SANS for selective observation of KaiA protomers/oligomers**

To clarify the kinetics of KaiA in a working Kai-Clock system, we conducted a time-resolved Contrast-Matching Small-Angle Neutron Scattering (tr-CM-SANS). Here, we explain the theory of CM-SANS and then also described our method for selective observation of KaiA protomers/oligomers in a working Kai-Clock system.

**SN2-1: Theory**

Neutron scattering has an isotope effect, which is largest in hydrogen atom: the scattering lengths of proton and deuteron are -3.74 fm and 6.671 fm, respectively. Therefore, by exchanging all hydrogen atoms with deuterium atoms in a protein, the scattering length density (SLD) of the protein is drastically increased. As shown in Fig.S9(a), the SLDs of hydrogenated (normal) protein (h-protein) and deuterated protein (d-protein) are 0.221 fm·Å$^{-1}$ and 0.774 fm·Å$^{-1}$, respectively: Here, it is reminded that there is different in SLD between amino acids but is almost no difference between proteins. SANS intensity of protein $I(\boldsymbol{q})$ is given as follows,

$$I(\boldsymbol{q}) = \int_V \left(\rho_p(\boldsymbol{r}) - \rho_s\right) \exp(-\imath \boldsymbol{q} \cdot \boldsymbol{r})\, d^3\boldsymbol{r} \qquad \text{(S1)}$$

$$|\boldsymbol{q}| = 2\left(\frac{2\pi}{\lambda}\right) \sin\left(\frac{\theta}{2}\right) \qquad \text{(S2)}$$

where $\boldsymbol{q}, \lambda$, and $\theta$ are scattering vector, wavelength of neutron, and scattering angle, respectively, and $\rho_p(\boldsymbol{r})$ is a distribution of SLD of protein as a function of potion $\boldsymbol{r}$**,** and $\rho_s$ is an averaged SLD of solvent. In equation (S1), the integral takes over the volume of protein $V$. Then, the forward scattering intensity $I(0)$ is also give as follows,

$$I(0) = \left(\overline{\rho_p} - \rho_s\right) \cdot V \qquad \text{(S3)}$$

where $\overline{\rho_p}$ is the averaged SLD of protein and $\left(\overline{\rho_p} - \rho_s\right)$ is called as "scattering contrast". As shown in Fig.S9(a), the SLD of 40% $D_2O$ solvent matches to that of (h-protein, meaning that the scattering intensity, especially $I(0)$, becomes zero and the h-protein is scatteringly invisible. Therefore, when h-protein and d-protein are mixed into 40% $D_2O$ solvent, the scattering of d-protein can be selectively observed (Fig.9(b)): This technique is named as Contrast-Matching SANS (CM-SANS).

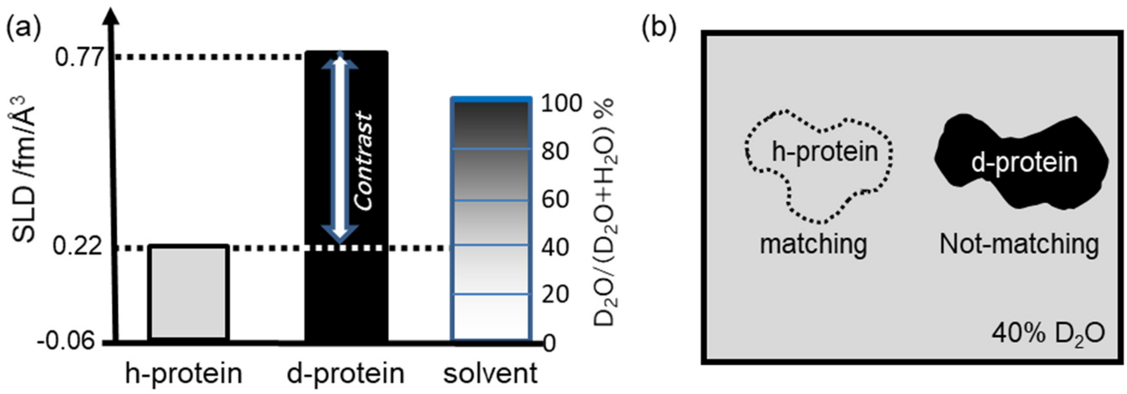


Figure S9. Contrast and CM-SANS. (a) SLDs of h-protein, d-protein and solvent with various $D_2O$ ratio. (b) Schematic illustration of CM-SANS technique. The h-protein and d-protein in 40% $D_2O$ solvent are scatteringly invisible and visible, respectively.

**SN2-2: Selective observation of KaiA in a working Kai-Clock system**

To selectively observe KaiA protein in a working Kai-Clock system which also includes KaiB and KaiC, we prepared for d-KaiA, h-KaiB and h-KaiC. In these three Kai proteins mixture solution with 40% $D_2O$ solvent, the only d-KaiA can be observed. Fig.S10 shows the schematic views of CM-SANS for the distributions of Points a-d as examples.

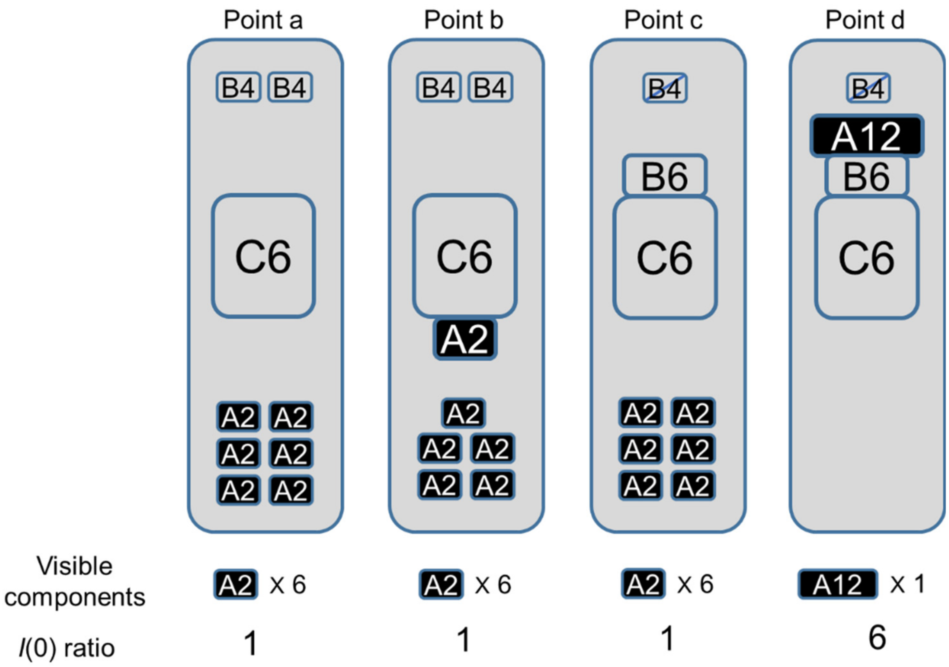


Figure S10. Schematic illustrations of CM-SANS for typical distributions. Six $A_2$ dimers are observed in Points a-c, whereas one $A_{12}$ is done in Point d, The $I(0)$ ratios is 1:6 for Points a-c and Point d.
